# The regulatory factor ELF1 triggers a critical wave of transcription in the antiviral response to type I interferon

**DOI:** 10.1101/549485

**Authors:** Leon Louis Seifert, Clara Si, Sarah Ballentine, Debjani Saha, Maren de Vries, Guojun Wang, Mohammad Sadic, Aaron Briley, Uwe Schäfer, Hong Moulton, Adolfo García-Sastre, Shashank Tripathi, Brad R. Rosenberg, Meike Dittmann

## Abstract

The transcription of interferon-stimulated genes (ISGs) is classically triggered via activation of the JAK-STAT pathway, and together, ISGs raise a multifaceted antiviral barrier. An increasing body of evidence reports the existence of additional, non-canonical pathways and transcription factors that coordinate ISG expression. Detailed knowledge of how heterogenous mechanisms regulate ISG expression is crucial for the rational design of drugs targeting the type I interferon response. Here, we characterize the first ETS transcription factor family member as a regulator of non-canonical ISG expression: E74-like ETS transcription factor 1 (ELF1). Using high-content microscopy to quantify viral infection over time, we found that ELF1, itself an ISG, inhibits eight diverse RNA and DNA viruses uniquely at multi-cycle replication. ELF1 did not regulate expression of type I or II interferons, and ELF1’s antiviral effect was not abolished by the absence of STAT1 or by inhibition of JAK phosphorylation. Accordingly, comparative expression analyses by RNAseq revealed that the ELF1 transcriptional program is distinct from, and delayed with respect to, the immediate interferon response. Finally, knockdown experiments demonstrated that ELF1 is a critical component of the antiviral interferon response in vitro and in vivo. Our findings reveal a previously overlooked mechanism of non-canonical ISG regulation that both amplifies and prolongs the initial interferon response by expressing broadly antiviral restriction factors.

**AUTHOR SUMMARY:** Over 60 years after their discovery, we still struggle to understand exactly how interferons inhibit viruses. Our gap in knowledge stems, on one hand, from the sheer number of interferon-stimulated effector genes, of which only few have been characterized in mechanistic detail. On the other hand, our knowledge of interferon-regulated gene transcription is constantly evolving. We know that different regulatory mechanisms greatly influence the quality, magnitude, and timing of interferon-stimulated gene expression, all of which may contribute to the antiviral mechanism of interferons. Deciphering these regulatory mechanisms is indispensable for understanding this critical first line of host defense, and for harnessing the power of interferons in novel antiviral therapies. Here, we report a novel mechanism of interferon-induced gene regulation by an interferon-stimulated gene, which, paradoxically, inhibits viruses in the absence of additional interferon signaling: E74-like ETS transcription factor 1 (ELF1) raises an unusually delayed antiviral program that potently restricts propagation of all viruses tested in our study. Reduced levels of ELF1 significantly diminished interferon-mediated host defenses against influenza A virus in vitro and in vivo, suggesting a critical but previously overlooked role in the type I interferon response. The transcriptional program raised by ELF1 is vast and comprises over 400 potentially antiviral genes, which are almost entirely distinct from those known to be induced by interferon. Taken together, our data provide evidence for a critical secondary wave of antiviral protection that adds both “quality” and “time” to the type I interferon response.

## INTRODUCTION

Within minutes of engaging their host cell receptors, type I interferons trigger a signaling cascade that results in the expression of hundreds of interferon-stimulated genes (ISGs) with antiviral activity (1). ISGs act on different stages of viral life cycles, from entry to viral genome replication, assembly, egress and finally, maturation (2). Given the plethora of diverse antiviral mechanisms, we are still striving to understand the complexity of how ISGs achieve their remarkably broad antiviral protection against positive, negative, and double-stranded RNA viruses, as well as DNA viruses and even intracellular bacteria and parasites (3–5).

The speed of the interferon signaling cascade is crucial for providing efficient antiviral protection, and is enabled by signaling components already present at baseline that eliminate the need for *de novo* protein synthesis (6). Three constitutively expressed transcription factors mediate interferon-stimulated gene expression: signal transducer and activator of transcription 1 and 2 (STAT1/2) (7) and interferon response factor 9 (IRF9) (8). Upon activation, phosphorylated STAT1/2 and IRF9 form the interferon-stimulated gene factor 3 (ISGF3) complex, which shuttles to the nucleus and initiates transcription from interferon-sensitive response elements (ISREs) (9). For over two decades, this has been our understanding of canonical type I interferon-signaling (10) (Fig. 1A, solid arrows).

**Fig. 1.**
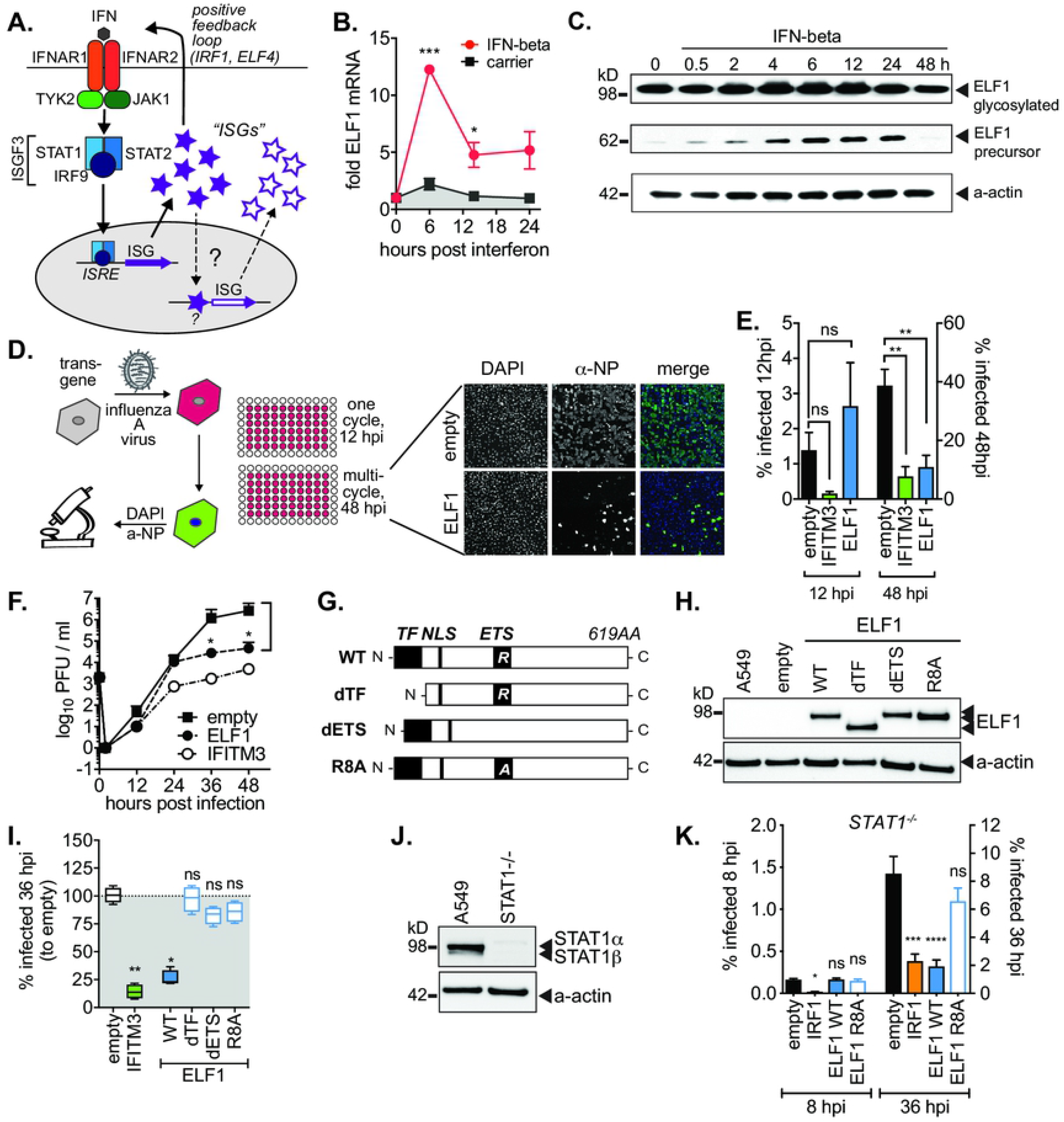
ELF1 is an interferon-stimulated transcription factor with delayed, STAT1-independent antiviral activity. (A) Canonical IFN signaling (solid arrows) and proposed second wave of ISG expression (dashed arrows). (B) Mean ± SEM of mRNA fold increase over pre-treatment control by qRT-PCR in A549 treated with IFN-beta or BSA carrier control (n=3). Paired t-test to BSA, *p<0.1, ***p<0.001. (C) Time course of ELF1 protein expression by western blot in A549 treated with IFN-beta. Two different exposures. ELF1 68 kDa, precursor ELF1; ELF1 98 kDa, glycosylated ELF1; a-actin loading control. (D) Mean ± SEM of % influenza A/WSN/1933 virus-infected (NP-positive) cells by high content microscopy in A549 expressing ELF1, IFITM3 as early (entry) ISG inhibitor control, or empty vector as negative control (n=3). 8 hpi (one cycle of replication, left y-axis) or 48 hpi (multi-cycle replication, right y-axis). One-way ANOVA and Dunnett’s multiple comparison test versus “empty”. (E) Representative images of one field of vision at 48 hpi from (C). (F) Influenza A/WSN/1933 virus growth kinetics on A549 expressing ELF1, IFITM3 or empty vector (n=3). Virus titers by plaque assay on MDCK cells. Individual t-tests to empty, *p<0.1, **p<0.01. (G) ELF1 protein domains and mutational strategy. TF, transcription factor domain; NLS, nuclear localization signal; ETS, E26 transformation-specific domain; R, conserved arginine within DNA binding domain; A, alanine substitution of conserved arginine within DNA binding domain. (H) Protein expression of mutant ELF1 by western blot. (I) Box and whiskers ± SEM of % influenza A/WSN/1933 virus-infected (NP-positive) cells by high content microscopy in A549 expressing ELF1, IFITM3 or empty vector (n=3). One-way ANOVA and Dunn’s multiple comparison test versus “empty”. *p<0.1, **p<0.01. (J) STAT1 protein expression in A549 STAT1−/− or control cells by western blot. K. Mean ± SEM of % influenza A/WSN/1933 virus-infected (NP-positive) cells by high content microscopy in STAT1−/− A549 expressing ELF1, ISG and transcription factor IRF1 as positive control, or empty vector (n=3). ANOVA with Dunnett’s multiple comparison to empty. **** p<0.0001, *** p<0.001 * p<0.01.

Recently, a number of studies have added astonishing complexity to the interferon signaling network by reporting multiple non-canonical mechanisms of ISG regulation, both dependent on and independent of JAK-STAT signaling (reviewed in (11)). While these important studies have begun to untangle the complexity of signaling mechanisms within the interferon response, and even led to the discovery of specific ISG subsets with antiviral activity, we are only starting to grasp that the interferon response also has a temporal gene expression component. Using novel transcriptional profiling techniques, several recent studies reveal that genes downstream of interferon signaling can be classified into qualitatively distinct modules with different temporal expression dynamics post-interferon (12–14). Such differences cannot be explained by the action of ISGF3 alone, suggesting non-canonical mechanisms at work (Fig. 1A, dashed arrows). How temporal divergence may influence the antiviral potency of the interferon response, and which factors or pathways drive divergent gene expression dynamics, remain unknown. However, an in-depth understanding of these factors and their specific contributions to the type I interferon response is crucial for defining the nature of host defenses and associated detrimental pro-inflammatory effects.

Despite evidence of temporal dynamics, most well-characterized ISGs act on early steps of viral life cycles. Hence, in search of “late-acting” ISGs, we previously performed a screen for ISGs inhibiting influenza A virus (IAV) specifically at multi-cycle viral replication. Intriguingly, we identified a transcription factor, E74-like ETS transcription factor 1 (ELF1), that fit this late-acting profile (15). Among hundreds of ISGs in our gain-of-function screen, expression of ELF1 alone was sufficient for potent virus inhibition with no detectable cytotoxicity. Its uniquely late antiviral action and antiviral potency positioned ELF1 as a putative novel regulator of a late module of antiviral genes. We speculate that ELF1’s exclusively late action could explain why it has not been previously identified as antiviral. Here, we characterize ELF1’s contribution to the type I interferon response, and show that it regulates a critical, previously unidentified set of potent antiviral restriction factors in a uniquely delayed fashion.

## RESULTS

### ELF1 is an interferon-stimulated gene with delayed, STAT1-independent antiviral activity

Previous studies on ELF1 focused on its role in lymphocyte maturation and immune signaling (16–18). To our knowledge, except for our ISG screen, ELF1 has never been characterized with regard to viral infections, and is completely uncharacterized in non-hematopoietic cells (15). Thus, we first validated ELF1 as an ISG in non-lymphoid cells. We used two airway epithelial cell culture systems relevant for IAV infection: A549, one of the most commonly used human epithelial cancer cell lines (Fig. 1B,C, and Fig. S1A), and stratified human airway epithelium cultures, a primary cell culture system that closely mimics the airways (19) (Fig. S1B). ELF1 mRNA and protein were expressed at baseline, and further induced by IFN-beta stimulation in both systems. Hence, ELF1 indeed acts as an ISG in airway epithelial cells.

Next, we confirmed ELF1’s delayed antiviral activity by high-content microscopy. This technique allowed us to monitor events occurring during viral infections at sub-cellular resolution. A549 cells were transduced to express ELF1 and controls, then challenged with a low MOI of IAV. IAV-infected cells were visualized at both 12 hpi (early stages of replication) and 48 hpi (late stages or multi-cycle replication) by immunostaining, and then quantified by high-content microscopy (Fig. 1D). As expected, we found that the ISG IFITM3, an early-acting positive control that blocks IAV fusion during entry, inhibited influenza A/WSN/1933 (H1N1) virus at 12 hpi, and continued to inhibit it at 48 hpi (Fig. 1E). In contrast, ELF1 did not inhibit IAV during single-cycle replication, but did inhibit multi-cycle replication (Fig. 1D,E), recapitulating the results from our screen.

In order to determine exactly when ELF1-mediated virus inhibition starts, we assessed low MOI multi-cycle growth kinetics in 12-hour increments on both A549 and primary normal human epithelial cells (NHBE). Viral titers from cells expressing exogenous ELF1 were significantly reduced for both cell types by at least 100-fold. While expression of IFITM3 decreased viral titers at 12 hpi, ELF1’s action was delayed, starting at 36 hpi (Fig. 1F and Fig. S2). Additional, detailed IAV life cycle studies revealed that this delayed antiviral action was not due to inhibition of individual IAV life cycle steps, such as entry, genome replication, egress or infectivity (Fig. S3A-E). Taken with its published role as a transcription factor, we hypothesized that ELF1 mediates its antiviral activity through the regulation of a transcriptional program that inhibits IAV at multiple levels during its replication cycle.

To test whether ELF1’s antiviral action is indeed through its activity as a transcription factor, we generated ELF1 mutants lacking ETS-transcription factor domains (Fig. 1G-H). These domains were all known or predicted by sequence homology with other related ETS transcription factors, such as ELF4. We deleted either the putative transcription factor (TF) domain, predicted to recruit RNA polymerase, or the ETS domain, which contains the DNA binding domain (16). Furthermore, within the ETS domain, we alanine-substituted an arginine (R8) that is conserved and critical for DNA binding in all ETS-transcription factors (20) (Fig. 1G). All three mutant proteins were expressed at similar levels as WT ELF1 (Fig. 1H). However, they lost their ability to inhibit IAV (Fig. 1I), supporting the hypothesis that ELF1 inhibits IAV through its transcription factor activity. From here on, we used the minimal ELF1 R8A mutant as a negative control for our study. Next, we assessed whether ELF1 executes its antiviral program through canonical interferon signaling. Theoretically, gene expression dynamics post-interferon could occur through a second round of canonical signaling in the form of positive feedback mechanisms, leading to temporal divergence (Fig. 1A, solid arrows). An example would be ELF4, which exerts its antiviral function through feeding-forward to produce more interferon (21). Another example of such a positive feedback loop is the ISG IRF1, which also triggers the production of interferon, as well as regulates its own set of antiviral ISGs (22). First, we tested by qRT-PCR whether ELF1 induces expression of type I or II interferons. In contrast to IRF1 control, which induced expression of interferon alpha, beta, kappa and gamma, ELF1 did not induce expression of any tested type I or II interferons (Fig. S4A). We thus reasoned that ELF1 might not exert its antiviral function through a positive feedback loop.

To examine this further, we performed ISRE reporter assays to test whether ELF1 induces gene expression from the ISRE element, the regulatory element recognized by ISGF3 (9). Other human ELF family members (2, 3, 4 and 5) were also tested; MDA5 served as positive, and GFP as negative control (Fig. S4B). We found that, in contrast to MDA5 and ELF4 (21), ELF1 does not induce transcription from the ISRE reporter.

Finally, we also repeated our IAV multi-cycle replication assay with ELF1 or positive control IRF1 on A549 lacking STAT1 (23) (Fig. 1J), a critical component of ISGF3 (7) (Fig. 1A). We found that although IRF1 induced expression of interferons (Fig. S4A), in accordance with its dual mechanism of antiviral action, it was still able to inhibit IAV in the absence of STAT1. ELF1 also potently inhibited IAV in the absence of STAT1. However, there was a striking difference in timing between the effects of IRF1 and ELF1: IRF1 inhibited IAV after one cycle of viral replication (as previously reported (4)), but ELF1 inhibited IAV exclusively in multi-cycle replication. Taking these results together, we hypothesized that ELF1 is produced in concert with other ISGs, then drives the expression of a distinct, delayed set of putative antiviral genes that without a second round of canonical interferon signaling (Fig. 1A).

### ELF1 regulates a vast transcriptional program, which is distinct from and delayed with respect to the immediate IFN response

Hence, we aimed to determine the transcriptional program regulated by ELF1 by performing RNAseq on A549 cells lacking ELF1. However, elevated amounts of ELF1 have been previously associated with lung cancer (24), suggesting that ELF1 may play a critical role in A549 growth. Accordingly, a viable clonal A549 ELF1^−/−^ knockout line could not be generated by CRISPR/Cas9 genome editing. To maintain a consistent cell type across experiments, we instead expressed ELF1 wild type (WT), R8A, or empty vector control in A549 (Fig. 2A, left column). Principal component analysis (PCA) indicated that WT ELF1 modulates gene expression patterns differently than ELF1 R8A and empty vector controls (Fig 2B). This analysis additionally validated ELF1 R8A as a negative control in our experiments.

**Figure 2.**
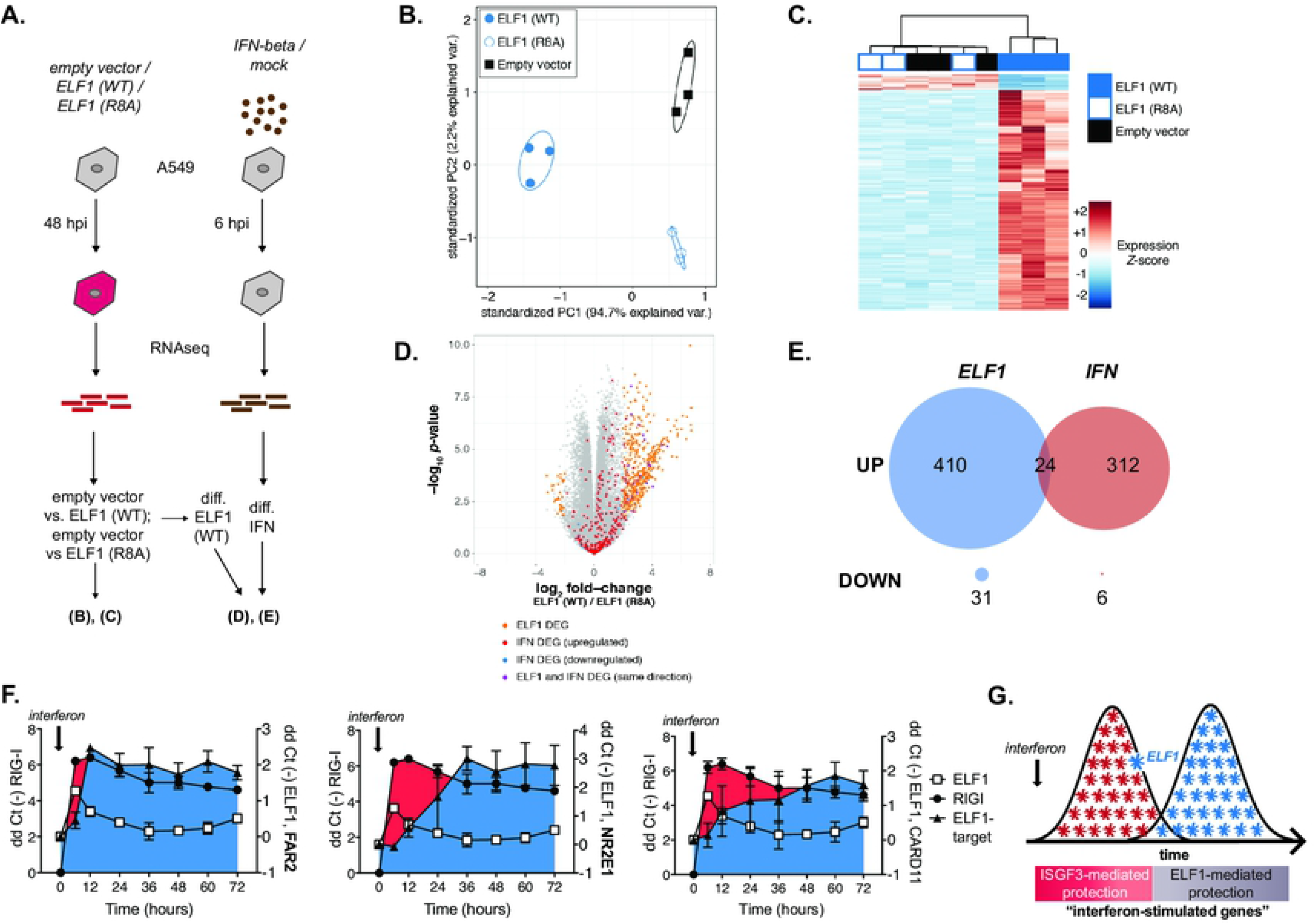
ELF1 regulates a vast transcriptional program, which is distinct from and delayed with respect to the immediate IFN response. (A) Schematic of RNAseq approach to identify genes expressed differentially in A549 expressing empty vector, ELF1 wild type (WT), or ELF1 loss-of-function mutant R8A, or in A549 treated with IFN-beta for 6 h. (B) Principal component analysis of 1000 most variable genes across all samples from empty vector, ELF1 wild type (WT), or ELF1 R8A-expressing cells (n=3, regularized log transformed counts, corrected for replicate batch effects). Ellipses indicate 68% normal probability for each group. (C) Heatmap plotting z-scaled expression values for ELF1 differentially expressed genes (n=3, adjusted p-value < 0.05, log2 fold-change >2 or <-2). (D) Volcano plot of WT ELF1 vs R8A ELF1 differential expression, also highlighting genes differentially expressed upon IFN stimulation. Each dot indicates one gene; and red dots expression upon ELF1 WT expression (n=3, adjusted p value < 0.05, log2 fold-change >2 or <-2), orange dots upon ELF1 expression and IFN-beta stimulation, and purple dots upon IFN-beta stimulation only. (E) Venn diagram depicting ELF1-vs IFN-beta-up or down-regulated genes. (F) Mean +/− SEM of mRNA expression (ddCT-) by qRT-PCR in A549 treated with IFN-beta. RIG-I (a prototype IFN-stimulated gene, left y-axis), ELF1 and ELF1 target genes FAR2 (left), NR2E1 (center) and CARD11 (right) on right y-axis, n=3. (G) Temporal schematic of ISGF3-mediated vs ELF1-mediated antiviral gene expression. Both pools of antiviral genes would be classified as “interferon-stimulated genes”.

We then performed differential gene expression analyses to identify genes for which expression changed upon ELF1 WT expression. Ectopic expression of ELF1 WT significantly altered the expression of 465 genes (Fig. 2C, Supplementary Tables 1, 2), most of which (434) were upregulated relative to both control conditions in additional comparisons. Analyzing the set of ELF1 differentially expressed genes, we made three key observations: 1. Gene ontology analyses of ELF1 differentially expressed genes revealed that many of the most significantly enriched terms relate to cell membrane and/or receptors (Fig. S5, Supplementary Table 5). 2. ELF1 does not trigger expression of IFN type I, II or III genes, or other inflammatory cytokines such as TNF or IL-6 (Fig. S6, Supplementary Tables 1 and 2). Indeed, IL-6 was downregulated upon ELF1 WT expression. These findings corroborated our previous results from qRT-PCR and ISRE reporter assays (Fig. S4), but were contrary to a previous study that found ELF1 to enhance the transcriptional response to IFN-beta (25). The differences might indicate cell-type specific differences between HeLa cells (25) and A549 cells in the present study. 3. Genes differentially expressed upon ELF1 WT expression were not enriched for GO terms implicated in IAV egress or infectivity, such as cargo trafficking, membrane remodeling, or maturation proteases, providing additional evidence that the ELF1-mediated antiviral program likely does not target specific steps of the IAV life cycle.

We next examined how different the ELF1 differentially expressed program was from the immediate IFN response. Historically, the term “interferon-stimulated gene” has been defined quite arbitrarily, depending on cell type, fold upregulation, and time post-interferon-stimulation (26). Typically, interferon-stimulated transcriptomes are determined experimentally only a few hours post-IFN exposure (4). To compare the ELF1 differentially expressed program to a typical ISG program, we treated A549 with IFN-beta for 6 h, performed RNAseq, identified immediate ISGs by differential expression testing (same statistical thresholds as above) and compared them to the differentially expressed genes in the ELF1 WT condition (Fig. 2A). Genes most dramatically upregulated (i.e. genes with highest-fold change) mostly did not overlap with ISGs rapidly induced by IFN stimulation (Fig 2D and Supplementary Tables 1, 2 and 3); of the 434 ELF1 differentially expressed genes relative to controls (by both adjusted p value and -fold change), only 24 cleared similar significance thresholds by IFN stimulation (Fig. 2E). Thus, although both the ELF1 differentially expressed program and the immediate IFN response program confer antiviral protection, they appear to be distinct, which leads us to one key conclusion: ELF1 differentially expressed genes represent a previously untapped pool of novel viral restriction factors.

To follow up on differences in timing of viral restriction, we determined the temporal dynamics of ELF1-associated gene transcription within the concert of the type I interferon response. Other studies have conducted similar experiments with endpoints up to 24 h post-interferon (12, 27). But, given ELF1’s delayed antiviral action, we followed mRNA expression of five indicator genes over a period of 72 h: our gene of interest, ELF1; a prototype of the immediate interferon-response, RIG-I; and three ELF1 differentially expressed genes that were amongst those genes most significantly upregulated upon ELF1 WT but not by 6 h IFN stimulation, FAR2, NR2E1 and CARD11 (Supplementary Tables 1, 2 and 3). As expected, RIG-I and ELF1 displayed expression kinetics typical for immediate ISGs, with peak expression occurring at 6 h post-IFN stimulation, followed by a gradual decrease to eventually reach homeostasis (Fig. 2F). In accordance with their presumed regulation by ELF1, upregulation of FAR2, NR2E1 and CARD11 temporarily succeeded that of ELF1 and RIG-I. Upregulation of FAR2 peaked at 12 h post-IFN stimulation, which was 6 h post-ELF1 peak expression. NR2E1- and CARD11-expression peaked even later, at 48 and 60 h post-IFN stimulation, respectively. These data demonstrate that putative ELF1 target genes are expressed with delayed kinetics relative to immediate ISGs, which might explain the observed delayed mode of IAV inhibition previously observed.

We thus propose the following temporal model of ELF1-mediated antiviral protection (Fig. 2G): upon interferon-stimulation, ELF1 is rapidly expressed in concert with hundreds of other immediate ISGs, in a process regulated by ISGF3. Subsequently, ELF1 acts as a direct regulator and initiates the expression of a distinct, second wave of novel antiviral genes. In this way, ELF1 both amplifies and lengthens the immediate antiviral interferon response via a novel transcriptional program. All of these genes qualify as “interferon-stimulated genes”, however, we show that their expression is based on different mechanisms of transcriptional regulation – i.e. the action of different transcription factors such as ELF1. These mechanistic differences have important consequences with regard to rational drug design to modulate the power of the interferon response—its activation for antiviral applications, and down-regulation for anti-inflammatory purposes.

### ELF1 inhibits multi-cycle replication of diverse RNA and DNA viruses

Thus far, we characterized the timing and composition of ELF1’s antiviral program. Next, we tested the breadth of ELF1’s antiviral action. We determined single and multi-cycle replication of eight diverse RNA and DNA viruses in the presence of ELF1 wild type and R8A (Fig. 3A): influenza A/WSN/1933 (H1N1) virus, human parainfluenzavirus 3 (HPIV3), yellow fever virus (YFV), chikungunya virus (CHIKV) (all three enveloped +RNA viruses), coxsackie B virus (CxB, a non-enveloped +RNA virus), herpes simplex virus 1 (HSV-1), vaccinia virus (VV) (both enveloped DNA viruses), and adenovirus 5 (AdV, a non-enveloped DNA virus). The ISG and transcription factor IRF1 again served as positive control, as it is known to inhibit all of these viruses in single cycle assays (3, 4). Our panel of viruses was chosen to broadly represent different viral mechanisms, e.g. how they enter cells and deliver their genomes, how they initiate viral transcription, how they replicate their genomes, how they process their proteins, how progeny viruses exit cells, how they counteract cellular immune responses, and more. These differences are reflected by the different replication rates of these viruses, which we considered when designing our single and multi-cycle assays (Fig. 3B-I, indicated at bottom of x-axes). We found that both IRF1 and ELF1 significantly inhibited all viruses in the panel (Fig. 3B-I), indicating that the breadth of ELF1-mediated virus inhibition is similar to that mediated by IRF1 and the immediate IFN response. However, and as seen in previous experiments, the difference between IRF1 and ELF1 was in timing, as ELF1 inhibited all viruses exclusively at multi-cycle replication (Fig. 3B-I). Interestingly, this multi-cycle antiviral action was apparent irrespective of the virus life cycle length. Therefore, it is possible that inherent differences in cells being challenged for the first time (single cycle virus infection) versus being challenged repeatedly (multi-cycle virus infection) contribute to ELF1’s delayed antiviral activity. Importantly, YFV and CHIKV are both extremely sensitive to endogenous interferon in A549 cells and were thus assayed in the presence of a JAK1/2 inhibitor, Ruxolitinib, to suppress JAK-STAT signaling and allow for viral replication (Fig. 3 D, E, and Fig. S7). ELF1 inhibited both YFV and CHIKV in the presence of Ruxolitinib. This validated our previous data from STAT1^−/−^ A549 cells (Fig. 1A), further supporting that ELF1 inhibits viruses without a second round of canonical interferon signaling.

**Figure 3.**
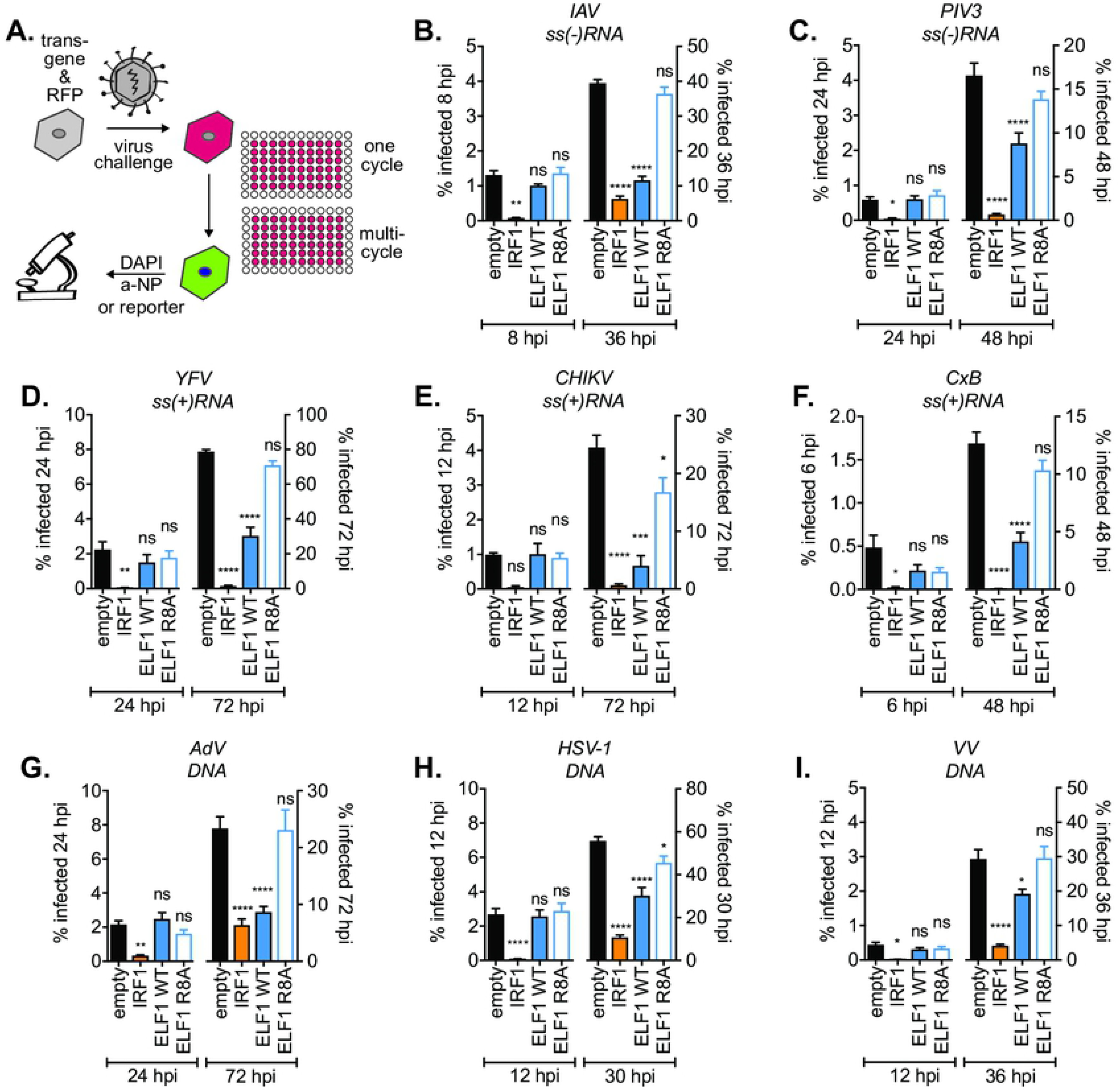
ELF1 inhibits multi-cycle replication of diverse RNA and DNA viruses. (A) A549 were transduced to express empty vector as negative control, ISG and transcription factor IRF1 as positive control, ELF1 wild type, or ELF1 R8A, a DNA binding domain mutant. 48 h post transduction, cells were challenged with a low MOI of the indicated viruses and % of infected cells determined by high content microscopy. Mean ± SEM of % virus-infected cells (n=3) at one replication cycle (left y-axes) or multi-cycle viral replication (right y-axes): (B) influenza A/WSN/1933 (H1N1), % NP-positive cells, (C) human parainfluenzavirus 3-EGFP, (D) yellow fever virus-Venus, (E) chikungunya-virus-ZsGreen, (F) coxsackievirus-EGFP, (G) adenovirus-EGFP, (H) herpes simplex virus 1-EGFP, or (I) vaccinia virus-EGFP. One-way ANOVA and Dunn’s multiple comparison test versus “empty” of the respective time point, *p<0.1, **p<0.01, ***p<0.001, ****p<0.0001.

### ELF1 is a critical component of the interferon response to influenza A virus in vitro and in vivo

How important is the ELF1-mediated gene expression module for the antiviral potency of the overall type I interferon response? To answer this question, we applied a knockdown strategy in A549. Treatment with peptide-conjugated phosphorodiamidate morpholino oligomers (PPMO) designed to bind the 5’UTR of ELF1 mRNA resulted in a 55% reduction of ELF1 protein (Fig. 4 A,B). We measured IAV growth in ELF1-PPMO-treated cells and found increased IAV titers over time (Fig. 4C). This increase started at 36 hpi and was most pronounced at 48 hpi, where viral titers were elevated up to 100-fold, demonstrating that endogenous ELF1 is a critical component of the antiviral interferon response *in vitro.*

**Figure 4.**
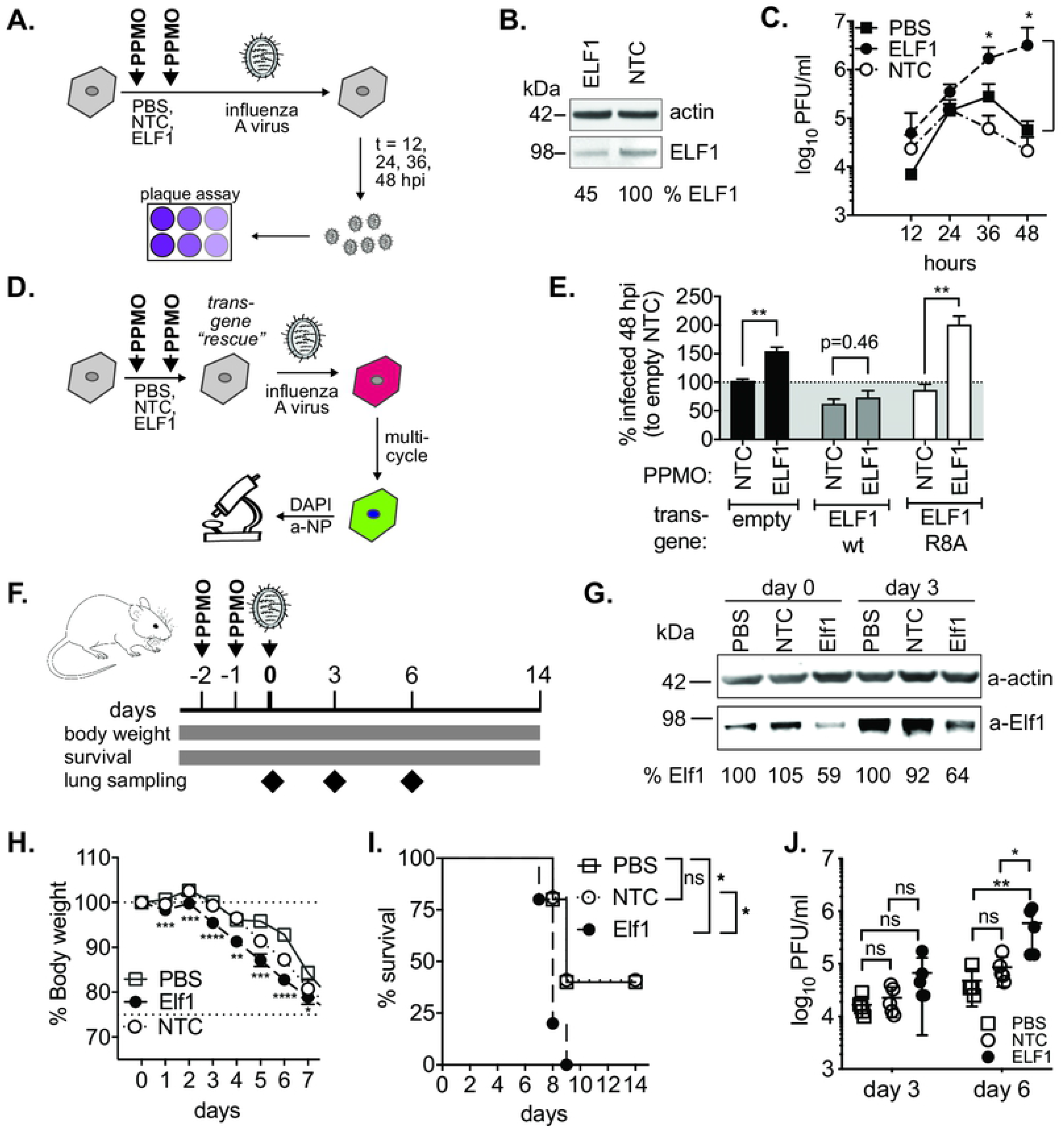
ELF1 is a critical component of the interferon response to influenza A virus in vitro and in vivo. (A) Schematic of peptide-conjugated morpholino (PPMO)-mediated ELF1 knockdown in A549 and influenza A/WSN/1933 virus low MOI growth kinetics. ELF1, ELF1 5’UTR-targeting PPMO; NTC, 5-base-pair non-targeting mismatch control. (B) Endogenous ELF1 protein and actin control post PPMO knockdown prior to infection (A) by western blot. % ELF1 protein normalized to actin and mismatch control. (C) Influenza A/WSN/1933 growth kinetics post PPMO knockdown, n=3. Mean ± SEM virus titer by plaque assay on MDCK cells. (D) Schematic of PPMO-mediated knockdown and transgene rescue in A549 expressing ELF1 wild type, R8A, or empty negative control. (E) Mean ± SEM of % influenza A/WSN/1933 virus-infected (NP-positive) cells by high-content microscopy, n=3. t-test comparing matching NTC and ELF1-knockdown samples, **p<0.01. (F) Schematic of PPMO-mediated in vivo knockdown. PPMOs targeting Elf1, a control mismatch PPMO, or PBS, were administered to BalbC mice intranasally at two and one days prior to infection. Mice were then challenged with 40 PFU of influenza A/PR8/1934 virus, and monitored for body weight and survival. Lungs were collected at days 0, 3 and 6 post infection for western blot analyses (G) and lung titer determination (J). (H) Mean ± SEM from PPMO-treated or control mice, n=15 mice per group. Unpaired two-tailed t-test comparing Elf1 to PBS, *p=0.1, ** p<0.01,***p<0.001, **** p<0.0001. (I) % survival (>25% body weight loss) of PPMO-treated or control mice, n=5 mice per group. Log-rank Mantel-Cox test, *p<0.1. (J) Virus titers in mouse lung homogenates were measured by plaque assay on MDCK cells, from n=5 mice per group. Mann-Whitney test, *p<0.1, **p=0.01.

To verify the specificity of PPMOs, we knocked down endogenous ELF1 in A549 with the ELF1-PPMO, then aimed to rescue ELF1’s antiviral function by providing ELF1 wild type as a transgene (Fig. 4D, E). As the ELF1-PPMO targets the 5’-UTR, it represses the translation of endogenous, but not overexpressed, ELF1. We used empty vector and ELF1 mutant R8A as negative controls, and visualized IAV infection at 48 hpi by high-content microscopy. As expected, knocking down endogenous ELF1 boosted the number of IAV-infected cells; neither empty vector control nor ELF1 R8A were able to rescue this phenotype (Fig. 4D, black and white bars). In contrast, expression of ELF1 wild type reduced IAV late stage or multi-cycle replication to levels that were similar in NTC and ELF1-PPMO-treated cells (Fig. 4D, grey bars). These results validated the specificity of our PPMO-mediated knockdown.

Finally, we sought to establish Elf1’s relevance *in vivo*, using the IAV infection mouse model. Previously generated and characterized Elf1^−/−^ mice were viable, but no longer available (http://www.informatics.jax.org/allele/MGI:3590647). Therefore, we induced a local knockdown by administering PPMO intranasally, as described previously (28). PPMOs targeting the 5’UTR of Elf1 and either a non-targeting PPMO mismatch control or PBS, both negative controls, were administered twice prior to IAV challenge (Fig. 4F). The Elf1-targeting PPMO yielded approximately 40% *in vivo* knockdown, as determined by quantitative western blot of mouse lung homogenates (Fig. 4G). PPMO-treated or control mice were infected intranasally with 40 PFU of influenza A/PR8/1934 (H1N1) virus. Animals with reduced Elf1 lost significantly more body weight and showed significantly increased mortality: 100% of Elf1-knockdown animals succumbed to infection, as compared to 50% in either control. Finally, Elf1-knockdown animals had significantly increased virus titers in the lung (Fig. 4H-J). Thus, Elf1 plays a pivotal role in the type I interferon response against IAV *in vivo.*

## DISCUSSION

In this study, we determine that ELF1 is an important antiviral regulator in vitro as well as in vivo, given that mice with decreased Elf1 protein levels exhibit more pronounced weight loss, higher mortality, and increased virus titers following influenza A virus challenge. Elf1’s *in vivo* role could either be fulfilled by cellular immunity (i.e. by lymphocytes), a cell-intrinsic function (i.e. in non-hematopoietic cells), or both. Indeed, lymphocyte functions of Elf1 have been previously published. The *in vitro* data presented in this study establish that ELF1 additionally exerts a cell-intrinsic antiviral role. Such different roles in specific cellular contexts are known for a number of other innate immune transcription factors. For example, the transcription factor IRF1 has been shown to both support the development of CD8+T and natural killer cells (29, 30), and to raise a cell-intrinsic antiviral program by activating IFN- and ISG-expression in fibroblasts, epithelial cells and skeletal muscle cells (31–34). Therefore, depending on the cellular context, it is feasible that ELF1 similarly displays dual functions.

ELF1 is an example of non-canonical interferon-mediated gene regulation that is independent of STAT1. ELF1 both adds “time” and “quality” to the immediate type I interferon response (Fig. 2H). These findings raise questions about the evolutionary rationale for multiple gene expression modules post-interferon. One explanation could be to provide multiple layers of mechanistically different, antiviral genes to cover antiviral strategies for as many diverse viruses as possible (4). Indeed, functional redundancy is a common theme of the innate immune response (35). Another rationale for switching antiviral programs could be to mitigate adverse effects brought about by prolonging existing ones. The multi-layered antiviral state raised by interferon represents a double-edged sword, as the associated inflammation can have detrimental effects (36). This is exemplified by genetic defects in key mediators of innate immune signaling that ultimately cause interferon overproduction or failure to return cells to homeostasis post-interferon exposure (36). Interestingly, multiple independent genome-wide association studies found single nucleotide polymorphisms in the ELF1 open reading frame or in ELF1 target DNA binding sites to be associated with chronic inflammatory disorders such as Crohn’s disease, inflammatory bowel disease, and systemic lupus erythematosus (37–43). How ELF1 may contribute to excess inflammation in these disorders remains elusive. However, transcriptional dysregulation, e.g. of IFN- or NFκB-mediated programs, has been shown to contribute to disease pathogenesis (44–49). These findings and ours raise the exciting possibility that ELF1 has a major regulatory function both in inflammation and in innate antiviral immunity.

Defining non-canonical interferon response programs such as that raised by ELF1 may pave new avenues in the rational drug design targeting the type I interferon response. Transcription factors have historically been considered to be undruggable, but this paradigm is slowly shifting (50). Novel small molecules are being developed that mimic DNA binding properties (51) or disrupt protein-protein interactions critical for transcription factor function (52), and artificial ligands aim to modulate transcription factor activation (53). There is an unmet need for novel antiviral drug targets, especially to combat emerging viruses, recently exemplified by Zika and Ebola viruses(54, 55). ELF1 inhibits every virus we have tested in this study, including members from diverse (-)RNA, (+)RNA, and DNA virus families. Thus, harnessing the antiviral power ELF1 might be an attractive approach for broadly antiviral therapies.

## MATERIALS AND METHODS

Detailed information on materials, including sources, are listed in Supplementary Table S6.

### Contact for reagent and resource sharing

Further information and requests for resources and reagents should be directed to and will be fulfilled by the Lead Contact, Meike Dittmann.

### Experimental model and subject details

#### Animals

Five-week-old female BALB/cJ mice were purchased from Jackson Laboratories (stock number 000651).

#### Ethics Statement

All research studies involving the use of animals were reviewed and approved by the Institutional Animal Care and Use Committees of the Icahn School of Medicine at Mount Sinai and were carried out in strict accordance with the recommendations in the Guide for the Care and Use of Laboratory Animals.

#### Primary human cells

Primary human normal airway tracheobronchial epithelial cells from de-identified donors (NHBE) (sex: male and female) were provided by Lonza (Walkersville, MD) and grown in BEM media supplemented with the BEGM bullet kit (Lonza). NHBE were used for functional studies, and for generation of polarized human airway epithelial cultures (HAE). To generate HAE, NHBE from individual donors were expanded on plastic to generate passage 1 cells, which were subsequently plated (5×10^4^ cells/well) on rat-tail collagen type 1-coated permeable transwell membrane supports (6.5mm; Corning Inc). HAE cultures were grown in B-ALI medium supplemented with inducer (Lonza Inc.) at each media change with provision of an air-liquid interface for approximately 6 weeks to form differentiated, polarized cultures that resemble *in vivo* pseudostratified mucociliary epithelium.

#### Cell lines

A549 (human; sex: male), A549 CRISPR STAT1^−/−^, HeLa (human; sex: female), HFF (human; sex: male), 293T (human; sex: female), 293T LentiX, 293T ISRE reporter, and MDCK (canine; sex female), LLC-MK2 (rhesus macaque), Vero (African green monkey; sex female) cells were maintained in DMEM (Invitrogen) supplemented with 10% fetal bovine serum (FBS), 1% NEAA, 1% P/S. A549 and HFF were used for virus infection and functional studies. MDCK, HeLa and LentiX cells were used for virus production and virus titration. All cell lines were obtained directly from the ATCC (with exceptions of Lenti-X 293T cells, which were obtained from Clontech Laboratories, and MDCK cells, which were obtained from the laboratory of Wendy Barclay). All cell lines were grown at 37 °C, individually expanded, and all seed and working stocks tested negative for contamination with mycoplasma. Cells were used in experiments below passage 15 from thaw, or when population doubling times slowed beyond 25% of seed stock doubling times.

#### Viruses

Influenza A/WSN/1933 (H1N1) virus stock was grown in MDCK cells. The following virus stocks were grown as previously described: HPIV3-GFP (based on strain JS) on LLC-MK2 cells (56), CxB-GFP (based on pMKS1-GFP) on HeLa cells (57), YFV-Venus (YF17D-5’C25Venus2AUbi) on Vero cells (58), HSV-1-GFP (based on strain Patton) on Vero cells (59), VV-GFP (derived on strain western reserve) on HeLa cells (Schoggins et al., 2011).

AdV-GFP (based on AdV5) was generated by the Laboratory of Patrick Hearing. The AdV5 E4-ORF3 reading frame was precisely replaced with EGFP in plasmid pTG3602 (60) using PCR, and recombineering in *E. coli*, as previously described (61). To generate infectious virus, the pTG3602-EGFP plasmid was linearized with PacI and 1 μg DNA transfected into 293T cells. Plaques were purified and working virus stocks were generated by passaging virus on 293T cells. The optimum dose for viral assays was determined by limited dilution and high content microscopy for EGFP-positive cells on A549.

Experiments with all above viruses were carried out in biosafety level 2 (BSL2) containment in compliance with institutional and federal guidelines. The infectious clone of CHIKV La Réunion 06-049 expressing ZsGreen was constructed by the Laboratory of Kenneth Stapleford using standard molecular biology techniques. First, an AvrII restriction enzyme site was inserted 5’ of the subgenomic promoter by site-directed mutagenesis using the primers Forward 5’-CACTAATCAGCTACACCTAGGATGGAGTTCATCCC-3’ and Reverse 5’-GGGATGAACTCCATCCTAGGTGTAGCTGATTAGTG-3’. The CHIKV subgenomic promoter was then amplified by PCR (Forward 5’-CCTAGGCCATGGCCACCTTTGCAAG-3’ and Reverse 5’-ACTAGTTGTAGCTGATTAGTGTTTAG-3’) and subcloned into the AvrII site to generate a CHIKV infectious clone containing two subgenomic promoters. Finally, the ZsGreen cassette was amplified by PCR (Forward 5’-GTGTACCTAGGATGGCCCAGTCCAAGCAC-3’ and Reverse 5’-GCTATCCTAGGTTAACTAGTGGGCAAGGC-3’) from a CHIKV infectious clone obtained from Andres Merits (University of Tartu) and subcloned into the AvrII restriction enzyme site. The complete cassette and subgenomic regions were sequenced to ensure there were no second-site mutations. To generate infectious virus, the plasmid was linearized overnight with NotI, phenol-chloroform extracted, ethanol precipitated, and used for in vitro transcription using the SP6 mMessage mMachine kit (Ambion). In vitro transcribed RNA was phenol-chloroform extracted, ethanol precipitated, aliquoted at 1 mg/ml, and stored at −80 °C. 10 μg of RNA was electroporated into BHK-21 cells (62) and virus was harvested 48 h post electroporation. Working virus stocks were generated by passaging virus over BHK-21 cells and viral titers were quantified by plaque assay. Experiments with CHIKV were carried out in biosafety level 3 (BSL3) containment in compliance with institutional and federal guidelines.

#### Lentiviral generation and transduction of cells

ISGs with antiviral activity (ELF1, IFITM3, IRF1, BST2) were part of the pSCRPSY lentiviral ISG library and co-expressed tagRFP and a puromycin resistance gene (Dittmann et al., 2015). To generate ELF1 domain deletion and point mutants, ELF1 wild type was amplified using forward primer 5’-ATGGCTGCTGTTGTCCAACAGAAC-3’ and reverse primer 5’-CTAAAAAGAGTTGGGTTCCAGCAGTTC-3’, and cloned into pCR8/GW/TOPO TA (Life Technologies). This entry clone DNA was used as starting point for mutagenesis. Mutation R8A was generated by site directed mutagenesis using Quikchange technology (Agilent), and forward primer 5’-TATGAGACCATGGGAGCAGCACTCAGGTACTATTAC-3’ and reverse primer 5’-GTAATAGTACCTGAGTGCTGCTCCCATGGTCTCATA-3’. ELF1 lacking the transcription factor (TF) domain was generated by PCR amplification of N-terminally truncated ELF1, using forward primer 5’-ATGGCTGCTGTTGTCCAACAGAAC-3’ and reverse primer 5’-CTAAAAAGAGTTGGGTTCCAGCAGTTC-3’ and cloning into pCR8/GW/TOPO TA. ELF1 lacking the internal ETS domain was generated using a PCR overlap extension PCR approach. We amplified the N-terminal fragment of ELF1 with forward primer 5’-ATGGCTGCTGTTGTCCAACAGAAC-3’ and reverse primer 5’-GGTGGATT CTAAAGCAGTGTCCAGGG CAAAAGTGGAAGGT CAG-3’, and the C-terminal fragment with forward primer 5’-GCAGTGTCCAGGGCAAAAGTGGAAGGTCAGCGCTTGGTGTATC-3’ and reverse primer 5’-CTAAAAAGAGTTGGGTTCCAGCAGTTC-3’. We then performed overlap extension PCR with the N-terminal and C-terminal PCR products as template, using forward primer 5’-ATGGCTGCTGTTGTCCAACAGAAC-3’ and reverse primer 5’-CTAAAAAGAGTTGGGTTCCAGCAGTTC-3’. The final PCR product was cloned into pCR8/GW/TOPO TA. From pCR8/GW/TOPO TA, ELF1 R8A, dTF and dETS constructs were swapped into pSCRPSY vector by gateway cloning.

To generate lentiviral stocks, we co-transfected 293T Lenti-X cells (Clontech laboratories) with plasmids expressing VSV-G, gag-pol, and the respective pSCRPSY plasmid, at a DNA ratio of 1:5:25. 48-72 h post transfection, we harvested the supernatant, centrifuged to remove cellular debris, and filtered the supernatant through a 0.2 μM filter. We then added HEPES to a final concentration of 20 mM and polybrene to 4 μg/ml. Each lentivirus stock was titrated on the respective cell types and diluted to obtain 90% transduced cells as determined by high-content microscopy.

#### IAV low MOI growth kinetics

To determine IAV growth, A549 or NHBE in 24-wells were transduced to express ELF1 and controls. 48 h post transduction, cells were gently washed twice with prewarmed PBS, and infected with influenza A/WSN/1933 (H1N1) virus at MOI 0.01 in 200 μl of PBS. The remaining inoculate was stored at −80 °C for back-titration. Cells with inoculate were placed in a rocking incubator at 37 °C for 1 h, then washed twice with prewarmed PBS, covered with 560 μl of prewarmed growth medium, and placed into a regular CO_2_ incubator at 37 °C. After 1 h, 50 μl of supernatant were collected and stored at −80 °C to determine successful removal of input virus. Supernatant was then collected every 12 h until 48 hpi, and stored at −80 °C. During supernatant collections, 50 μl of fresh, prewarmed growth medium were replaced in each well to keep total volume constant throughout the 48 hours.

#### IAV plaque assay

IAV infectious titers were determined by plaque assay on MDCK cells. Briefly, MDKC cells in 12-well plates were washed with PBS, and incubated with 1:10 serial dilutions of IAV in PBS for 1 h in a rocking incubator. After 1 h, cells were washed and overlayed with DMEM (Gibco), 1.2 % avicel, 0.001 % DEAE, 0.45 % Sodium Bicarbonate, GlutaMax, non-essential amino acids, penicillin/streptomycin, and 1 μg/ml TPCK trypsin. Cells were then placed in a CO2 incubator at 37 °C for 48 h. To fix cells and visualize plaques, the avicel overlay was aspirated, washed once with PBS, and cells covered with 0.1 % crystal violet, 2 % ethanol, 20 % methanol for 15 min, then washed with water, and plaques counted manually.

#### High-content microscopy and image analysis

If not otherwise stated, we used the CellInsight CX7 High-Content Screening (HCS) Platform (Thermofisher) and high-content software (HCS) for microscopy and image analysis. All small molecule inhibitors used in this study were tested for cytotoxicity and optimum effective dose for each cell type. Cells in 96-wells were incubated with a serial dilution of the inhibitor, keeping the carrier concentration constant in each well. Incubation time corresponded to the time the drug would be in contact with the cells in the actual assay. 10 % Ethanol was used as positive control for cell death. Cells were then stained with Sytox green (1:20,000, Thermo), washed with DMEM, fixed with 1.5 % paraformaldehyde, permeabilized with 0.1% triton X-100, stained with DAPI, and imaged with the 4x objective. For this assay, we used the HCS analysis protocol “Target Activation”, and reference levels were set at three standard deviations from the mean of control wells. Cytotoxicity was evaluated by a reduction of total (DAPI-positive) cells per well, as well as the % of dead (Sytox-positive) cells. Drugs were used at the highest safe dose in the assays described below, i.e. the dose not reducing the number total cells, and not increasing the number of dead cells as compared to carrier control.

For virus spread experiments, the optimum MOI and timing of endpoints was determined by high content microscopy prior to experiments for each cell type and each virus. For experiments with the endpoint at one round of viral replication, we chose the time that resulted in bright, yet individual virus-positive cells. For the optimum MOI, we chose the viral dose that yielded reproducible 0.5-3 % infected cells at that time point. For the second endpoint at multiple rounds of replication, we chose the time that resulted in 10-60 % of infected cells from that viral dose, depending on the spreading capability of the given virus. Experiments with YFV and CHIKV were performed in the presence of 0.4 μM of Ruxolitinib to allow for viral spread on IFN-competent A549. Further experiments analyzing the action of antiviral ISGs (ELF1, IFITM3, BST2) were performed using these optimized viral doses and time points: A549 (or A549 STAT1^−/−^) in multiple 96-well plates were transduced with pSCRPSY:empty lentivirus for 48h, and then infected with serial dilutions of the respective viruses. At different times post-infection, each plate was fixed with 1.5 % paraformaldehyde for 15 min, washed with PBS, quenched for 5 min with 20 mM NH_4_Cl, and washed with PBS again. To permeabilize the cells, we used 0.1 % Triton-X in PBS for 4 min, followed by washing with PBS three times. For reporter viruses expressing a strong GFP-signal (HPIV3, YFV, CHIKV, AdV, HSV-1, CxB and VV), cells were stained with DAPI only. IAV-infected cells were blocked with 1% BSA in PBS for 1 h at room temperature, stained with anti-NP antibody (1:500, BEI resources) for 1 h rocking at 37°C, washed three times with PBS before staining with secondary goat Alexa 488 antibody and DAPI for 1 h rocking at 37°C and finally washed three times with PBS. Plates were imaged using the 4x objective in 9 fields covering the entire 96-well. For this assay, we used the HCS analysis protocol “Target Activation”, and reference levels were set at three standard deviations for the highest background from all mock-infected control wells.

#### ISG induction assays

To determine ISG mRNA expression and protein kinetics, we treated PBMCs, A549 cells or HAE cultures with IFN-beta (Millipore Sigma) at 500 U/ml. mRNA levels, normalized to housekeeping gene RPS-11, were determined by qRT-PCR (SuperScript III First Strand Synthesis System, Life Technologies and PowerUP SYBR Green Master Mix, Thermo Fisher Scientific). Primer sequences are listed in Supplementary table 9. Endogenous ELF1 protein levels from cell lysates or mouse lung homogenates were measured by western blotting using anti-ELF1 antibody (1:5000, Santa Cruz).

#### PPMO-mediate ELF1 knockdown in vitro

All PPMOs were tested for cytotoxicity and knockdown efficiency in A549 cells prior to infection experiments. PPMOs were delivered to cells by adding them to the medium. For IAV growth kinetics, A549 were supplemented with 15 μM PPMOs for 2 d, then infected with influenza A/WSN/1933 virus at MOI 0.01 as described above. During supernatant collections, 50 μl of fresh, prewarmed growth medium with 8 μM PPMOs were replaced in each well to keep total volume constant throughout the 48 h. For determination of IAV spread by high content-microscopy, A549 were transduced to express ELF1 wild type, ELF1 R8A, or empty vector control for 6h, then media was changed to media containing 15 μM PPMOs. After 2 d, cells were infected with 100 PFU/well of influenza A/WSN/1933 virus, and infection media supplemented with 8 μM PPMOs. Samples were fixed and analyzed at 8 hpi and 36 hpi as described above.

#### PPMO in vivo assays

Five-week-old female BALB/c mice were anesthetized by intraperitoneal injection of a mixture of Ketamine and Xylazine (100 μg and 5 μg per gram of body weight), prior to intranasal administration of either PBS or 100 micro moles of PPMO mix (50 micro moles of PPMO1 and 2 each) in 40 μl of PBS, on Day −2 and Day −1. On Day 0, Mice were challenged intranasally with 40 PFU of PR8 IAV (LD50 = 50 PFU) in 40 μl PBS. Mice were monitored daily for weight loss and clinical signs. Mice lungs were harvested on Day 3 and Day 6 post infection for measuring viral titers (5 mice per condition). Lung homogenates were prepared using a FastPrep24 system (MP Biomedicals). After addition of 800 μl of PBS containing 0.3% BSA, lungs were subjected to two rounds of mechanical treatment for 10 s each at 6.5 m/s. Tissue debris was removed by low-speed centrifugation, and virus titers in supernatants were determined by plaque assay. A group of mice (5 per condition) until day 14 post infection for survival.

#### RNA-seq and analysis

For ectopic ELF1 RNAseq experiments, A549 cells in 6-well plates were transduced with pSCRPSY lentivirus encoding ELF1 wildtype, ELF1 R8A, or empty vector control. After 48 h in culture, medium was aspirated, cells were washed with PBS, and cells were scraped from the plate and homogenized in 1 ml of Trizol. Cell lysates were transferred to phasemaker tubes (Invitrogen), to which 200 μl chloroform was added, followed by vigorous mixing for 20 s. After incubation at room temperature for 2-3 min, samples were centrifuged at 12,000xg for 10 min at 4 °C. 600 μl of the aqueous (top) phase was transferred to 600 μl ethanol and mixed well by vortexing. The aqueous phase/ethanol mixes were then transferred to RNeasy columns and RNA extracted following the RNeasy kit protocol (Qiagen). The experiment was performed three times, with each transduction condition represented once in each replicate “batch.” RNAseq libraries (for all samples in a single batch) were prepared with the Illumina TruSeq Stranded Total RNA Library Prep Kit according to manufacturer’s instructions, and sequenced on the Illumina NextSeq 500 platform at 75nt read length in single-end configuration.

Reads were mapped to the human genome reference (hg19) supplemented with the pSCRPSY plasmid sequence (containing EGFP gene), using the HISAT2 (v2.0.4) alignment tool (63) with Ensembl v75 gene annotations (supplemented with pSCRPSY gene annotation) and the “--rna-strandness R” and “--dta” parameters. Read counts per gene were quantified against Ensembl (v75) transcript reference annotations (appended with gene annotation for pSCRPSY, “MSTRG.1”) using HTSeq-count (v0.6.1p1) (64). Genes with greater than 2 read counts in at least 3 samples were defined as “expressed” and included in downstream analyses. For principal component analysis (PCA), read counts were normalized and variance stabilized by regularized log transformation (rlog function, DESeq2 package v1.18.1 (65)). Replicate batch effects were corrected with the removeBatchEffect function in the limma package. PCA was conducted on the 1000 most variable genes across all samples.

For differential gene expression analysis, raw read counts were TMM-normalized and log_2_ transformed with voom (limma v3.34.9) (66, 67). Differential gene expression testing was performed with a linear model including factors for ELF1 (WT, R8A, or empty vector) and replicate batch. Pairwise tests were conducted for ELF1 (WT) vs (ELF1 R8A), and ELF1 (WT) vs Empty vector contrasts. Differential gene expression test p values were adjusted for multiple testing by the method of Benjamini and Hochberg (http://www.jstor.org/stable/2346101). In order to focus further analyses on those genes markedly affected by ELF1, a relatively stringent filter was applied to differential expression results: “ELF1 differentially expressed genes” were defined as those genes with adjusted p value < 0.05 and log_2_ fold-change ≥ 2 (or < −2) in both ELF1 (WT) vs (ELF1 R8A), and ELF1 (WT) vs Empty vector contrasts. GO term enrichment analysis in ELF1 differentially expressed genes was performed with the GOSeq tool (v1.3) (68), and results visualized with GOplot (v1.0.2)(69).

For IFN-stimulation experiments, A549 cells were stimulated with IFN-beta at 500 U/ml for 6 h, or left untreated. RNA was extracted and prepared for RNA-Seq as described above. Libraries were sequenced on the Illumina HiSeq 2500 instrument at 50nt read length in single-end configuration. Reads were processed and quantified against Ensembl (v75) transcript reference annotations as for ELF1 transduction experiments above, with identical expression filters applied. Differential gene expression testing was performed using limma, with model including factors for interferon stimulation (IFN-stimulated or mock-treated) and replicate batch. Pairwise tests were conducted for the IFN-stimulated vs mock-treated contrast. As for the ELF1 transduction analysis, “IFN differentially expressed genes” were defined as those with adjusted p value < 0.05 and log_2_ fold-change ≥ 2 (or ≤ −2).

### Quantification and statistical analysis

All n of in vitro experiments are from biologically independent experiments. Statistical analysis was performed in Prism (GraphPad Software, v7.0f, 2018). The statistical tests used and the number of biological replicates is indicated in each figure legend. Unless otherwise stated two conditions were compared using two-tailed Student’s t-tests. Statistical significance was defined as a p value of 0.05.

### Data and software availability

All RNA-Seq data are available in NCBI GEO repository, combined in SuperSeries GSE122252.

## ACKNOWLEDGEMENTS

We would like to thank Charles M. Rice for mentorship and support to initiate this project, and for YFV-Venus; Victor Torres for PBMC; Shoshanna Kahne and Nathan Mercado for technical assistance; Peter Palese for all IAV isolates and Hans-Heinrich Hoffmann for generating virus stocks, Paula Traktman for VV-EGFP, Peter Collins for HPIV3-EGFP, Ann Palmenberg for CxB-EGFP, Kenneth Stapleford for CHIKV-ZsGreen, Ian Mohr for HSV-1-EGFP, and Patrick Hearing for AdV-EGFP. This work was supported by National Institutes of Health (NIH) grants K99/R00-AI121473, R01-AI091707, DP5OD012142, the Boehringer Ingelheim Foundation, NYU Medical School startup funds, and anonymous donations. This work was also partly supported by NIAID grant U19AI135972, and by CRIP (Center for Research on Influenza Pathogenesis), an NIAID funded Center of Excellence for Influenza Research and Surveillance (CEIRS, contract number HHSN272201400008C).

## AUTHOR CONTRIBUTIONS

M.D. designed the project and wrote the manuscript. M.D., S.T., A.G.-S., U.S. and B.R.R. conceived and designed the experiments. L.S., C.S., S.B., A.B., M.d.V, M.S., D.S., G.W., S.T., B.R.R., and M.D. performed the experimental work and analyzed the results. H.M. designed and synthesized PPMOs. All authors discussed the results and commented on the final manuscript.

## SUPPLEMENTARY INFORMATION

Supplemental Information includes seven figures and seven tables and can be found with this article online.

### Supplementary Figure legends

**Supplemental Fig. S1, related to Fig. 1 B,C.**

(A) A549 were treated with IFN-beta or BSA carrier control, and mRNA expression determined over time by qRT-PCR. Fold increase over pre-treatment control from n=3 independent experiments. Data for ELF1 and RSAD2 as immediate ISG control are shown side-by-side as as mean ±SEM. Paired t-test of each time point compared to carrier, *p<0.1, ***p<0.001. (B) H&E stain of human airway epithelial cultures. Human airway epithelial cultures were basolaterally stimulated with IFN-beta, and mRNA expression determined over time as described in (A). Data for ELF1 and RSAD2 shown as mean ±SEM of 3 technical replicates. Paired t-test of each time point compared to t=0, *p<0.1, **p<0.01, ***p<0.001.

**Supplementary Fig. S2, related to Fig. 1F.**

Normal human bronchiolar epithelial (NHBE) cells were were transduced to express the indicated ISGs and infected with influenza A/WSN/1933 virus at MOI 0.01. Virus titers in the supernatants were measured in the inoculate (0 h), after inoculation and washing (2 h), and in 12 h intervals, by plaque assay on MDCK cells. Individual t-tests compared to empty control, **p<0.01.

**Supplementary Fig. S4, related to Fig. 1 J,K.**

(A) A549 were transduced to express ELF1 wild type (wt), ELF1 loss-of function mutant R8A, IRF1 as positive control, or empty vector control. Expression of type I and II IFNs by qRT-PCR. Data is shown normalized to empty control, as mean ± SEM. **** p<0.0001, ** p<0.01. (B) Reporter assay for ISRE-driven transcription. 293T carrying firefly luciferase under the control of a promoter carrying the ISRE motif and stably expressing renilla luciferase as control was transfected to express GFP as negative control, MDA5 as positive control, or ELF1, ELF2, ELF3, ELF4 and ELF5, respectively. Cells were subsequently stimulated by transfection of polyI:C. Data as mean ± SEM from n=3 independent experiments.

**Supplementary Fig. S2, related to Fig. 1. ELF1 does not inhibit specific life cycle steps, but inhibits multi-cycle IAV replication.**

(A-E) A549 cells were transduced to express the indicated ISGs. Empty vector served as negative control, and the following positive controls were used for individual IAV life cycle steps: Diphyllin for IAV entry, Ribavirin for IAV replication, Oseltamivir for IAV budding and detachment, IFITM3 for IAV entry, BST2 for IAV egress. Data are represented as mean ± SEM from at least n=3 independent experiments for all panels. (A) A549 were challenged with influenza A/WSN/33 virus at MOI 1, and the number of NPpositive nuclei was determined by high-content microscopy at 6 hpi. 1-way ANOVA and Dunn’s multiple comparison test. *p<0.1, **p<0.01, ***p<0.001. (B) IAV replication efficiency was assayed by a luciferase-based IAV minigenome assay in 293T cells. Expression constructs for components of the IAV replication machinery (PB1, PB2, PA and NP, of A/WSN/1933 origin) were co-transfected with a reporter construct mimicking the viral genome, leading to expression of firefly luciferase when the genome mimic is replicated. Individual t-tests compared to empty control, ***p<0.001. (C) Influenza A/PR/8/1934-NS1-GFP virus single cycle replication was assayed by flow cytometry, determining the percentage of infected (GFP-positive) A549 at 10 hpi, in the ISG-expressing (RFP-positive) population. Individual t-tests compared to empty control, **p<0.01, ***p<0.001. (D.-E) A549 were infected with influenza A/WSN/1933 virus at MOI 1, washed, and assayed at 12 hpi. (D) viral RNA (vRNA) was extracted from supernatants, and vRNA copy number was determined by qRT-PCR. (E) Infectious virus titers in the supernatant were determined by plaque assay on MDCK cells. Individual t-tests compared to empty control, *p<0.1, **p<0.01, ***p<0.001.

**Supplementary Fig. S5, related to Fig. 2C.**

“Bubble plot” depicting GO terms enriched in ELF1 differentially expressed genes from Figure 2C. Each bubble represents a significant (GOSeq adjusted p-value < 0.05) GO term. y-axis indicates enrichment significance (-log10 adjusted p-value) and x-axis indicates gene expression fold-change score ([upregulated genes – downregulated genes]/ √number of genes]) for term member genes. Bubble size is proportional to the number of term member genes. GO categories (Biological Process, Cellular Component, Molecular Function) are presented as separate panels to facilitate visualization. Highly significant enriched GO terms (adjusted p-value < 10-5) are annotated.

**Supplementary Fig. S6, related to Fig. 2 C.**

A549 were transduced to express ELF1 wild type (wt), ELF1 loss of function mutant R8A, or empty vector control. 48 h post transduction, cells were harvested and mRNA expression determined by RNAseq. All data shown from n=3 biologically independent experiments. RNAseq read counts per million (not normalized) for type I and II IFNs, TNF and IL-6 (Ensembl v75) from (A) for each sample. Although positive read counts were detected for IFNE in all samples, IFNE was not differentially expressed in any contrasts across conditions. IL-6 was downregulated in ELF1 WT samples.

**Supplementary Fig. S7, related to Fig. 3. D and E.**

A549 cells were treated with indicated amounts of the pan-Jak inhibitor Ruxolitinib (Rux), or DMSO carrier control, and infected with YFV-Venus. Cells were imaged and cell numbers or YFV-Venus positive cells determined by high-content microscopy. (A) Cell count per well (DAPI-positive) 72 post Rux treatment. (B) % YFV-Venus positive cells. (C) Representative images of (A) and (B). (D) Analysis of STAT3 phosphorylation as a readout of Jak activity. A549 cells were treated with 500 U/ml of IFN-beta, indicated amounts of Rux, or DMSO carrier control. At 48h post treatment, cells were harvested and analyzed by western blotting using anti-pSTAT3 antibody, or anti-actin antibody as loading control.

### Supplemental Tables

Supplemental Table 1. RNAseq ELF1-wt vs empty, related to Figure 2.

Supplemental Table 2. RNAseq ELF1-wt vs R8A, related to Figure 2.

Supplemental Table 3. RNAseq IFN vs control, related to Figure 2.

Supplemental Table 4. Gene Ontology ELF1 all categories, related to Figure 2.

Supplemental Table 5. RNAseq 434 ELF1-wt unique genes, related to Figure 2.

Supplemental Table 6. Material and sources, related to Methods.

Supplemental Table 7. Oligonucleotides, related to Methods.

